# Soil viral and prokaryotic communities shifted significantly after wildfire in chaparral and woodland habitats

**DOI:** 10.1101/2024.11.15.623834

**Authors:** Sara E. Geonczy, Anneliek M. ter Horst, Joanne B. Emerson

## Abstract

Increased wildfire activity warrants more research into fire-driven biotic changes in soil, given that soil microbes contribute to biogeochemical processes by way of organic matter decomposition, nutrient cycling, and promoting plant growth. Viruses of prokaryotes apply pressure to microbial communities, making their responses to fire also important for understanding post-fire ecology. Leveraging viromes and 16S rRNA gene amplicon sequencing, here we studied viral and prokaryotic community responses to wildfire in woodland and chaparral soils at five timepoints over one year following the California LNU Complex wildfire.

We also compared post-fire samples to unburned controls at the final three timepoints, beginning five months after the fire. Viromic DNA yields were low-to-undetectable (indicative of low viral particle abundances), particularly for the first post-fire timepoint, and comparisons to controls suggest a return to baseline viral particle abundances within five months of the wildfire. Viral and prokaryotic community composition and soil chemistry differed significantly in burned samples compared to controls from both habitats. Compared to controls, a greater proportion of viral ‘species’ (vOTUs) from a burned conifer forest were detected in both burned habitats here, suggesting fire-associated habitat filtering. Published viromes collected from the same sites nine months pre-fire were more similar to controls than to post-fire viromes. Together, these results indicate significant changes in soil viral and prokaryotic communities due to wildfire.

## INTRODUCTION

The current increase in global fire activity, driven by climate change, land use change, and fire suppression policies, warrants a better understanding of fire’s effects on biogeochemical processes and ecosystem resiliency in Mediterranean shrubland and woodland habitats [1–3]. This includes the impact on prokaryotic communities, which are significant drivers of soil biogeochemical processes, with major roles in organic matter decomposition and nutrient cycling [4,5]. Changes in soil properties, which often occur after a disturbance like a wildfire [6], are well known to drive bacterial community shifts [7]. One indirect impact of fire is a decrease in soil moisture, which can impact microbial communities, especially in habitats with low precipitation already [8]. Soil viruses are also known to be impacted by drought and heat [9,10]. What are not so well known are the short- and long-term impacts of wildfire on soil viral communities and the roles that viruses may play in the post-fire environment.

Wildfires can transform landscapes and sometimes act as an ecosystem “reset”, driving primary and secondary plant succession [11]. Some habitats are wildfire mediated, including chaparral, with vegetation that evolved with fire that has adaptations both conducive and sometimes dependent on a certain frequency of fire activity [12]. Soil bacterial community shifts can influence ecosystem successional rates [13], which makes post-fire soil microbial dynamics of great interest, given the threat of loss of habitat after wildfires [13,14]. Studies with high-resolution temporal sampling after fire have shown rapid bacterial community change, including both dramatic shifts immediately after and rampant community compositional changes over time [12]. Pyrophilous bacteria and fungi have been identified for their proliferation after fire, which has been attributed to traits that confer success in a post-fire environment, such as heat tolerance, xerotolerance, and spore and sclerotia formation [15,16]. Additionally, pyrophilous bacteria seem to be phylogenetically conserved, which may mean they are both adapted to and ubiquitous across habitats that have experienced a fire disturbance [17]. Although heat-responsive vOTUs have been identified [9], the extent to which there might be ‘pyrophilous’ viruses is unknown.

While bacterial succession after wildfire has been studied in chaparral [12], little is known about the communities in woodland habitats that are often found close to chaparral. Additionally, there is little known about the viruses in these same systems in general [18], especially with regard to the post-fire response. Here we present a one-year temporal study on viral and prokaryotic communities in woodland and chaparral soils following a wildfire. Our aim was to compare responses between the two habitats over time while also comparing community responses to dynamics in non-fire-impacted controls and to viromes previously collected from the same sites nine months before the fire [18]. Additionally, we sought to place the viruses detected in disturbed and undisturbed samples in biogeographic context, assessing where they had been detected previously and whether the detection patterns might suggest conserved viral adaptations to fire.

## MATERIALS AND METHODS

### Study site description

The LNU Lightning Complex fires burned 147,000 ha in northern California and were active for 46 days from 17 August 2020 to 2 October 2020 [19]. On 4 November 2020 (approximately one month post-fire), we set up four 5×5 m plots at Quail Ridge Natural Reserve, CA, which burned at low-to-moderate severity in all locations throughout the field site, determined via the Monitoring Trends in Burn Severity (MTBS) dataset [20]. Two plots were in burned chaparral habitats and two in burned woodland habitats. Plots were spatially paired, meaning that each chapparal plot was located close to (within 85 m of) a spatially paired woodland plot. These plots were all considered burned plots. On 12 March 2021, we set up an additional four burned plots (two chaparral and two woodland, which were spatially paired within 150 m of each other) at McLaughlin Natural Reserve, CA and four control plots (two chaparral and two woodland, which were not spatially paired due to the constraint of sampling locations that had not been burned by the wildfire). We also leveraged seven viromes from seven sites collected in November 2019 as part of a separate study [18], allowing us to make pre-burn comparisons for the Quail Ridge site, which burned completely in the LNU Complex fires.

Overall, there were 19 sampling locations (**Supplementary Table 1**). Chaparral sites had loamy soil with minimal horizon development, with surface mineral soil (A horizon) extending down to 5 to 15 cm. Woodland sites had predominantly loamy clay-enriched soil with an A horizon down to 25 cm. There was no organic horizon at any of the sites [21]. Chamise was the dominant vegetation for all chaparral sites, and oak and pine trees were the dominant vegetation for woodland sites.

### Soil sampling

We collected two replicates per time point in each plot by dividing the plot in half along the middle, then dividing the subplot into eight sampling units, and collecting soil (0-6 cm depth) in corresponding sampling units at each timepoint (e.g., the top right of the right subplot and the top right of the left subplot for a single timepoint). The distance between replicate samples within each plot was approximately 3.75 m. We homogenized and sieved (8 mm) all samples separately. Samples were stored on ice and transported back to the lab and promptly placed at 4°C until DNA extraction two to three days after sample collection. We sampled five timepoints at Quail Ridge and three timepoints at McLaughlin (corresponding with the final three timepoints at Quail Ridge), with all timepoints two to three months apart, yielding a total of 88 samples.

### Viromic DNA extraction and quantification

Within two or three days after sample collection (12 samples were randomly processed each day), we extracted viromic DNA from all samples, as previously described [9,22]. Briefly, to 10 g of each soil sample, we added 9 mL of protein-supplemented phosphate-buffered saline solution (PPBS: 2% bovine serum albumin, 10% phosphate-buffered saline, 1% potassium citrate, and 150 mM MgSO4, pH 6.5). Samples were vortexed until homogenized and then placed on a horizontal shaker at 300 RPM at 4°C for 10 min. Supernatant was transferred into a new tube, and then 9 mL of PPBS was added to the soil remaining in the original tube, repeating the process for a total of three soil resuspensions and elutions, ending with a total of three volumes of supernatant pooled together in a new centrifuge tube. The pooled supernatant was then centrifuged at 8 min at 7840 RPM at 4°C, collecting the progressively clearer supernatant and discarding the concentrated soil particles, for a total of three centrifugations and supernatant collections. The final supernatant was filtered through a 5 μm polyethersulfone (PES) membrane filter (PALL, Port Washington, NY, USA) to filter out larger soil particles, and then the filtrate went through a 0.22 μm PES membrane filter (PALL) to exclude most cellular organisms (apart from ultra-small bacteria [10,23]). Viral-sized particles and DNA remaining in the filtrate were concentrated into a pellet using the Optima LE-80K ultracentrifuge with a 50.2 Ti rotor (Beckman-Coulter, Brea, CA, USA) at 32,000 RPM at 4°C for 2 h 25 min. The pellet was resuspended in 200 μL of 0.02 μm filtered ultrapure water and treated with DNase (briefly, incubation at 37°C for 30 min after adding 10 units of RQ1 RNase-free DNase, Promega, Madison, WI, USA). We then extracted DNA using the DNeasy PowerSoil Pro kit (Qiagen, Hilden, Germany), following the manufacturer’s protocol, with the addition of a 10 min incubation at 65°C prior to cell lysis, which also served as the ‘stop’ incubation for the DNase treatment.

DNA quantification was performed using the Qubit dsDNA HS Assay and Qubit 4 fluorometer (ThermoFisher, Waltham, MA, USA). Due to low (30 viromes) or undetectable DNA yields (below 0.1 ng/μl or 0.49 ng/g soil) (19 viromes) now known to be expected for dry, heated, and recently burned soils [9,10], a subset of 39 viromes was prepared for library construction and sequencing.

### Total DNA extraction and quantification

For each sample, 0.25 g of soil was added directly to the DNeasy PowerSoil Pro kit (Qiagen) for total DNA extraction, following the manufacturer’s protocol, with the addition of a 10 min incubation at 65°C prior to cell lysis. DNA quantification was performed using the Qubit dsDNA HS Assay and Qubit 4 fluorometer (ThermoFisher).

### Virome library preparation and sequencing

Sequencing of all 39 viromes was performed by the DNA Technologies and Expression Analysis Core at the University of California, Davis Genome Center (Davis, CA, USA). Metagenomic libraries were constructed with the DNA Hyper Prep kit (Kapa Biosystems-Roche, Basel, Switzerland) and sequenced (paired-end, 150 bp) using the NovaSeq S4 platform (Illumina, San Diego, CA, USA) to a requested depth of 10 Gbp per virome.

### Amplicon library preparation and sequencing

We performed 16S rRNA gene amplicon sequencing on all total DNA extractions with a dual-indexing strategy [24]. To amplify the V4 region of the 16S rRNA gene, we used the Platinum Hot Start PCR Master Mix (ThermoFisher) with the 515F/806R universal primer set. We followed the Earth Microbiome Project’s PCR protocol [25], which included an initial denaturation step at 94ºC for 3 min, 35 cycles of 94ºC for 45 s, 50ºC for 60 s, and 72ºC for 90 s, and a final extension step at 72ºC for 10 min. We cleaned libraries with AmpureXP magnetic beads (Beckman-Coulter), quantified amplicons with a Qubit 4 fluorometer (ThermoFisher), and then pooled all bead-purified products together in equimolar concentrations. The libraries were submitted to the DNA Technologies and Expression Analysis Core at the University of California, Davis Genome Center and sequenced (paired-end, 250 bp) using the MiSeq platform (Illumina).

### Soil physicochemical measurements

During sample collection, in-field soil temperature measurements were taken at each sampling location, and water repellency was measured by wetting a sample with a single drop of deionized water and counting how many seconds it took until the drop had fully infiltrated into the soil [26] (**Supplementary Table 2**). For the remaining soil physicochemical measurements, we note that there were some methodological differences among time points, which we clarify in detail here. In particular, differences between T0 (from our prior work at the same site) and T1-T5 are in part on account of not anticipating a need to compare these datasets, as well as differences in protocols before T0 and during (T1-T5) the COVID-19 pandemic. Differences within the T1-T5 dataset were largely for cost-effectiveness, as we learned in real time about costs and options at different testing facilities. For T1-5, we calculated gravimetric soil moisture of each sample, by weighing out 10 g of soil into an aluminum weigh dish and measuring the weight after 24 hours in a drying oven at 105°C. For T0, gravimetric soil moisture was calculated by weighing out 10 g of soil and allowing it to dry completely in a biological hood for up to several days before weighing again. For T0 (from our prior study), soil samples were sent to Ward Laboratories (Kearney, NE, USA) for basic soil tests. For T1, we sent a soil sample from each sample site (at this point, only the 8 samples from Quail Ridge) to A&L Western Agricultural Laboratories (Modesto, CA, USA) for basic soil chemistry analysis as well as the University of California Davis Analytical Lab (Davis, CA, USA) for additional soil chemistry analysis. For timepoints T2-T5, we homogenized subplot replicate soil samples and sent a representative 200 g of field-condition soil (not dried) to Ward Laboratories for a suite of soil chemistry analyses under the Haney Soil Health Analysis package [27,28]. Analysis was done by using the soil properties that we have available for all timepoints, which includes: gravimetric soil moisture (difference in method described above, soil pH (1:1 soil/water suspension), organic matter (percentage weight loss-on-ignition), nitrate (T0-1 was KCl-extracted and T2-5 was Haney-extracted), inorganic phosphorus (Olsen method for T0-1 and Haney-extracted and quantified by flow injection analysis for T2-5), and potassium, calcium, magnesium, and sodium (T0-1 was ammonium-acetate extracted and T2-5 was Haney-extracted). All Haney-extracted nutrients were quantified using inductively coupled argon plasma (ICAP) atomic emission spectroscopy (**Supplementary Table 3**).

### Virome bioinformatic processing

Raw reads were trimmed using Trimmomatic v0.39 [29] with a minimum q-score of 30 and a minimum read length of 50 bases. We used BBMap v39.01 [30] to remove PhiX sequences. Each virome was assembled separately with MEGAHIT v1.2.9 [31] in the metalarge mode with a minimum contig length of 10,000 bp. Assembled contigs were analyzed with VIBRANT v1.2.0 [32] with the -virome flag to identify viral contigs. All output viral contigs (low, medium, and high quality) from all samples were iteratively clustered to form a representative set of vOTUs. We used CD-HIT v4.8.1 [33] to cluster viral contigs at 95% shared nucleotide identity and 85% alignment coverage (breadth). Quality-filtered raw reads from all viromes were then mapped against the representative set of vOTUs using Bowtie 2 v2.4.1 [34] in sensitive mode. Lastly, we used CoverM v0.5.0 [35] to quantify vOTU relative abundances in each virome and generated a trimmed mean coverage table and a count table for downstream analyses (0.75 minimum covered fraction). Host taxonomy of all vOTUs was predicted using iPHoP v1.1.0 [36], with default parameters (minimum cutoff score of 90) (**Supplementary Table 4**).

### Virome read mapping to habitat-labeled vOTUs from other datasets

To investigate whether vOTUs in our dataset were previously found in other habitats, we mapped reads (with Bowtie 2 v2.4.1, as described before) to the PIGEON v2.0 vOTU database [37,38], which contained 515,763 vOTUs, and two combined datasets collected from Blodgett Research Forest [9,39], which contained 272,017 vOTUs. For the Blodgett datasets, we identified vOTUs that were only found in burned forest samples (labeled as burned mixed conifer forest), vOTUs that were only found in unburned forest samples (labeled as undisturbed mixed conifer forest), and vOTUs that were found in both burned and unburned forest samples (labeled simply as mixed conifer forest). We then retrieved sequence data for all vOTUs that met minimum coverage (CoverM – 0.75 minimum covered fraction) by using the filterbyname tool from BBMap v39.01 on both the PIGEON v2.0 database and the Blodgett Forest combined datasets. We determined the proportion of vOTUs found in our dataset that were previously found in each of the other habitats. Finally, we clustered these vOTUs with the rest of the dataset at 95% average nucleotide identity (ANI) using CD-HIT [33], redid read mapping to this updated set of vOTUs, and generated a new trimmed mean coverage table for downstream analysis.

### 16S rRNA gene amplicon sequence bioinformatic processing

We used DADA2 v1.12.1 [40] to demultiplex 16S rRNA gene amplicon reads and perform quality assessment, read merging, and chimera removal. For quality filtering, we used the filterAndTrim() function with the following parameters: truncLen = c(0,0), maxN = 0, maxEE = c(2,2), truncQ = 2, rm.phix = TRUE. All other processing was with default settings. Amplicon sequence variants (ASVs) were assigned taxonomy by using the DADA2 RDP classifier, using the SILVA database v138.1 as reference [41].

### Statistical and ecological analyses

R v4.4.1 was used to perform all ecological and statistical analyses [42]. Input for all analyses of viral communities or vOTUs was the trimmed mean coverage vOTU table. We normalized the vOTU abundance table by using the decostand function (method = “total”, MARGIN = 2) from the Vegan package v2.6-6.1 [43] and removed singletons, (vOTUs detected in only one sample). With a DADA2-generated abundance table for 16S rRNA gene amplified single variants (ASVs), we filtered out mitochondria and chloroplasts and removed singletons (ASVs that only appeared in a single sample). We calculated Bray-Curtis dissimilarities with vegdist (method = “bray”), performed PERMANOVAS with adonis2, and performed PERMDISPs with betadisper and permutest, all functions from the Vegan package. We performed multidimensional scaling and retrieved true eigenvalues to calculate PCoA points with the cmdscale function (base stats). To perform PCA analysis, we used the princomp function (base R). Upset pots were created with the ComplexUpset v1.3.3 package using species abundance tables that were transformed into presence-absence tables [44]. We determined distance between plots with the geosphere v1.5-18 package [45]. Statistical analyses are documented in **Supplementary Tables 5-7**.

## RESULTS AND DISCUSSION

### Study design and dataset features

The intention for this study was to assess time-resolved changes in dsDNA viral and prokaryotic communities after a wildfire, comparing the response in two habitats (chaparral and woodland) to control (not burned) plots. We set up a total of 12 5×5 m plots, 8 burned and 4 control, with half of the plots in chaparral habitat and the other half in woodland habitat (**Figure 1A**). Over the course of 10 months (or one year since the start of the wildfire), we sampled five timepoints (T1-T5) at Quail Ridge Natural Reserve (CA) and three timepoints at McLaughlin Natural Reserve (CA) (corresponding with T3-T5 at Quail Ridge) for a total of 88 samples. For comparison, we also leveraged a pre-burn dataset of seven viromes from the same field sites collected in November 2019 (9 months pre-burn) [18].

**Figure 1.**
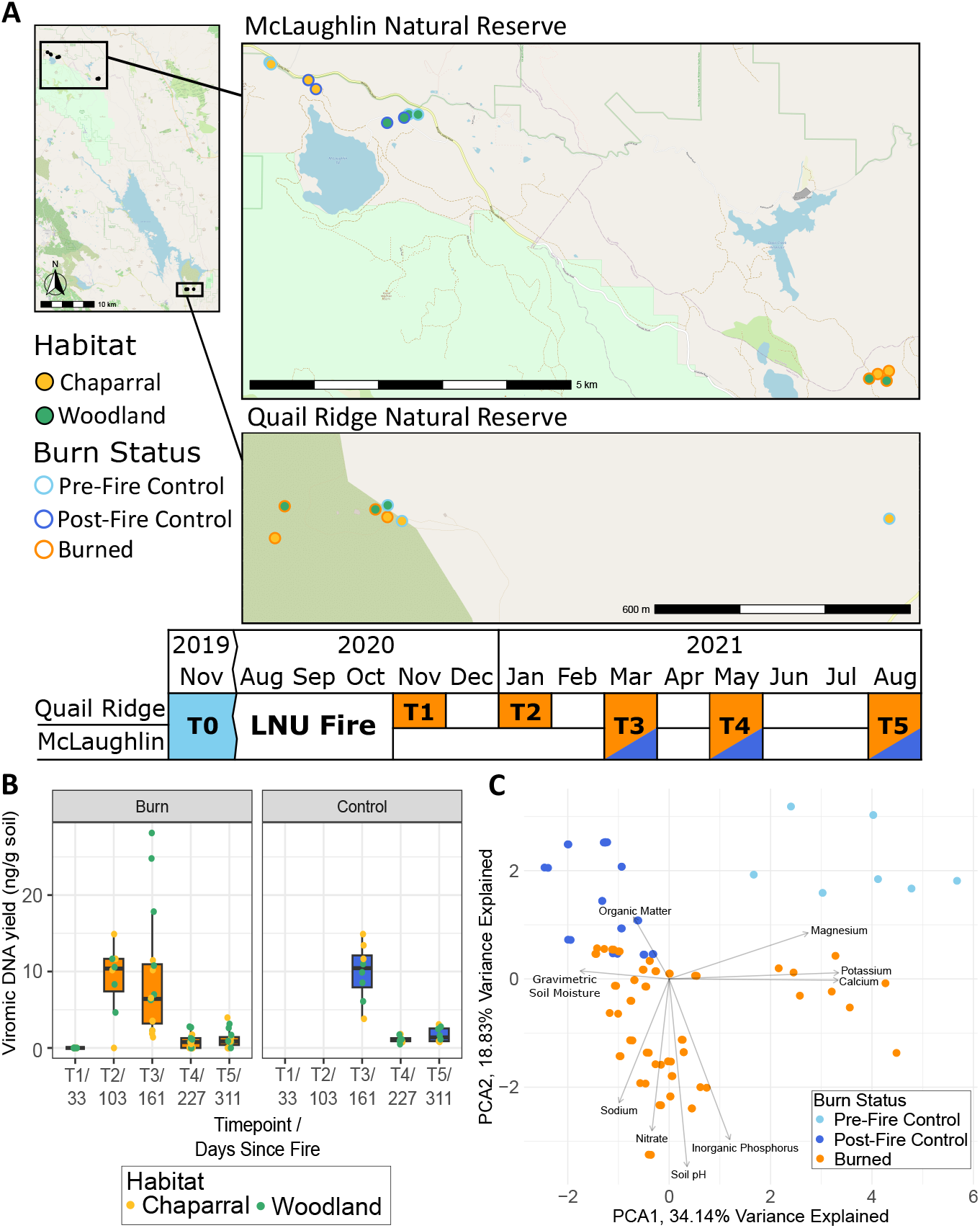
**A**. Map of plot locations in the main map and point fill indicating habitat and point outline indicating burn status in the subset maps, for a total of 12 plots. Two replicate cores were collected from each plot (each point in the map) at each sampled time point (except for the pre-fire controls). Table shows timeline of sample collection in each natural reserve. **B**. DNA yields for all viromes over time (labels show time point / days since fire), separated by burned (left facet) and control (right facet) plots. Dot color indicates habitat. Box boundaries correspond to 25^th^ and 75^th^ percentiles, and whiskers extend to ±1.5x the interquartile range. T1 and T2 were not collected for control plots but remain as “blanks” in the graph for easier comparisons to burned plots T3-T5. T0 (not plotted) was 322 days before the fire. **C**. Principal component analysis (PCA) of z-transformed physicochemical properties for each sample, with color indicating burn status.

We initially generated 88 DNase-treated viromes and used the viromic DNA yields to estimate patterns of viral particle abundances (as in prior work, [10]) and to determine which viromes to sequence (**Figure 1B**). Yields were below detection limits for all samples at T1 (all of which were burned samples), indicating low viral particle abundances [10], consistent both with the dry season at the time of sampling and the potential long-term temperature and moisture impacts of the fire [46,47], all of which have been shown to reduce viromic DNA yields [9,10].

For the three time points with control samples (T3-T5), there was no significant difference in viromic DNA yields between burned and control samples at any of the time points (Kruskal-Wallis, T3: P = 0.2446, T4: P = 0.2534, T5: P = 0.1581), suggesting a return to baseline viral particle abundances within 5 months of the fire (the first control timepoint). By T4 and T5, viromic DNA yields had decreased significantly in both chaparral and woodland habitats in both control and burned samples (Kruskal-Wallis, P < 0.05 for all conditions), presumably reflecting the transition into the dry season in the Mediterranean climate, which has previously been shown to substantially reduce the abundance of free viral particles [10]. Due to undetectable and low DNA yields, we elected not to sequence 49 of the viromes, which, to preserve statistical power for comparisons, reduced the number of time points considered for our viral community analysis. We sequenced 39 viromes, including all viromes from Quail Ridge (burned) from T2-T4 except one (Chaparral, T2) and all viromes from McLaughlin T3, which included the control viromes. We also extracted total DNA from all 88 samples and amplified the 16S rRNA gene region to profile prokaryotic communities. All 88 samples amplified and were sequenced, but one sample yielded insufficient sequencing data (Quail Ridge, Chaparral, T5), so it was removed from the analysis during rarefaction. We recovered 5,097 ASVs over the entire dataset (**Supplementary Table 8**), which reduced to 4,933 ASVs after rarefying to a common depth of 17,700 reads per sample and removing singletons.

In our bioinformatic analyses, we also included seven viromes that were collected from chaparral and woodland habitats at both Quail Ridge and McLaughlin in November 2019 [18], nine months prior to the fire. Paired 16 rRNA gene amplicon profiles were planned for that dataset, but, due to the COVID shutdown, they were never generated, and the soils are no longer available. The viromes served as pre-fire controls and increased the number of vOTUs used for read mapping-based vOTU detection in our dataset. In total, we amassed 77,869 putative viral contigs > 10 kbp as identified by VIBRANT [32], 77,433 of which were de novo assembled from this study, and 436 of which were vOTUs from the 2019 viromes. We also mapped reads to vOTUs from the PIGEON v2.0 database [38] and from viromes generated from our recent prescribed burn study in a forest [39], resulting in detection of 1,644 additional vOTUs. All recovered viral contigs and vOTUs were then clustered into a dereplicated set of 71,046 vOTUs (**Supplementary Table 9**), which was reduced to 37,926 vOTUs after removing singletons (vOTUs detected in only one sample).

### Viral communities differed significantly in burned and unburned samples and between habitats, whereas fire homogenized prokaryotic communities across habitats

Since soil chemical properties are a good indicator of fire impact [6], we first compared the profile of chemical properties across soil samples. A principal component analysis revealed significant separation of burned samples from post-fire controls based on soil chemistry (p = 0.04 by PERMANOVA) (**Figure 1C**). The loadings in the direction of the burned samples relate to post-fire responses reported elsewhere, including increased pH [48] and increased available phosphorus [6,11]. We then compared beta-diversity of viral and prokaryotic communities in post-fire and control samples to see whether they differed significantly. Burn condition (burned or unburned) significantly structured viral communities (explaining 13.1% of the variation, p < 0.001 by PERMANOVA) (**Figure 2A**). Habitat explained 5.82% of the variation (p < 0.001 by PERMANOVA), still playing a part in viral community structuring, as has been observed previously [18]. Despite large spatial differences in sampling locations, spatial distance (plot location) did not seem to obscure treatment effects on viral communities as substantially as has been observed in other studies [22,49], including in our analysis of viral community responses to a prescribed burn in a California mixed conifer forest [39] (p < 0.001 by PERMANOVA for plot location across all viromes, and p < 0.001 across all prokaryotic profiles). This means that plots with the same condition (burned or unburned) or the same habitat were most similar, overcoming plot-specific variability. While we did see viral community compositional structuring by habitat in control samples, as in prior work [18], habitat had a more pronounced influence on viral community composition post-fire (interaction of habitat and burn condition explained 8.1% of the variation in the data, P < 0.001 by PERMANOVA). The dispersion of viral communities was significantly greater in burned viromes than in controls (p = 0.006 by PERMDISP), meaning that burned viral community composition had greater variability than in unburned controls. When considering vOTU detection patterns in each habitat, 30% of the 37,926 vOTUs were detected only in post-fire burned woodland and 13% only in post-fire control woodland. For chaparral, 23% were found only in post-fire burned locations and 11% only in post-fire control locations.

**Figure 2.**
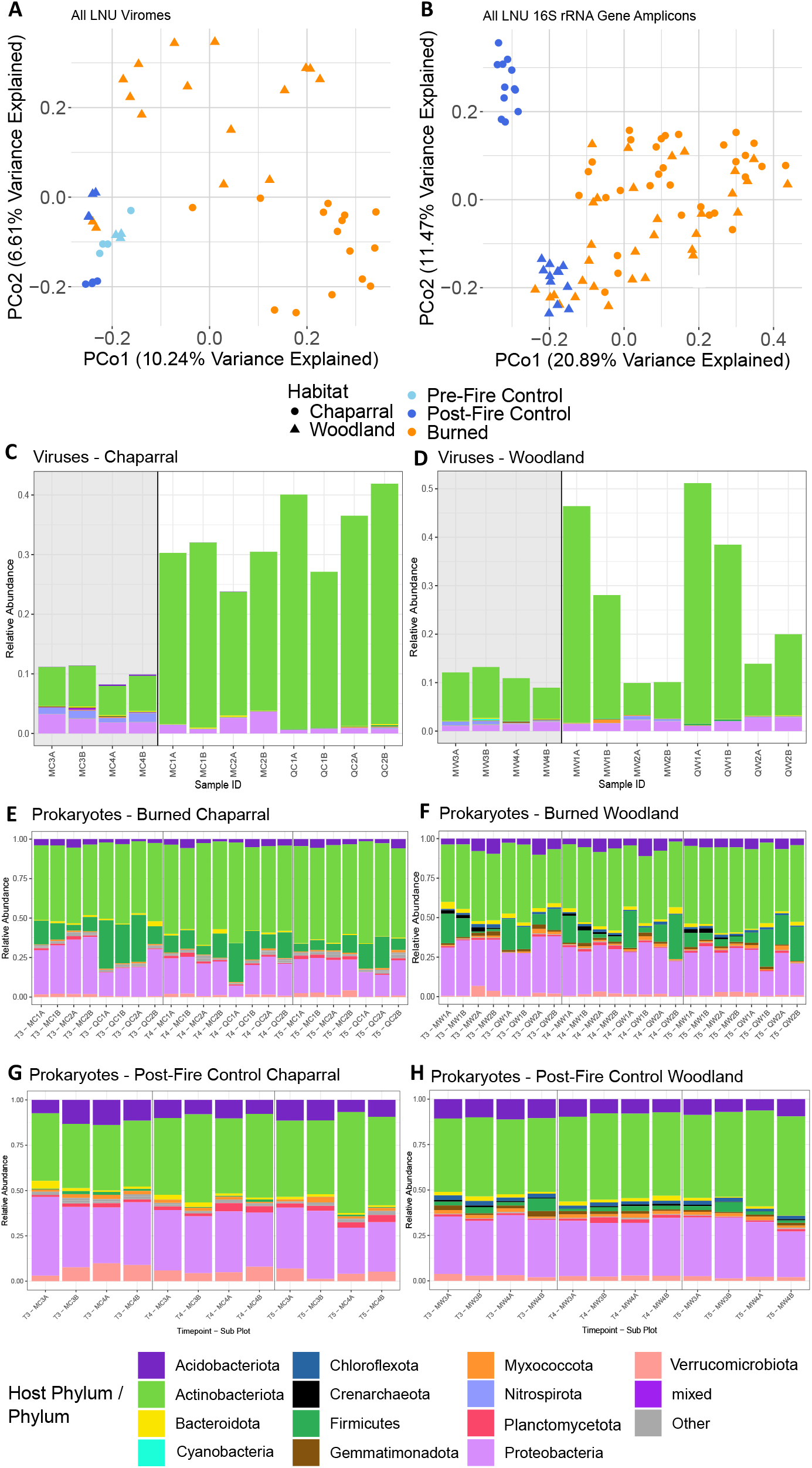
**A/B**. Unconstrained analyses of principal coordinates (PCoA) performed on vOTU (**A**) and prokaryotic ASV (**B**) Bray-Curtis dissimilarities calculated across all viromes or ASV gene profiles (from 16S rRNA gene amplicon data), respectively, with color indicating burn condition and shape indicating habitat. **C/D**. Relative abundances of vOTUs labeled with their phylum-level host predictions for chaparral (**C**) and woodland (**D**) habitats. Each stacked bar is one virome. Less abundant host phyla are collapsed into the “other” group. Panel background colors highlight control (gray) and burn (white) samples. **E-H**. Phylum-level relative abundances in 16S rRNA gene profiles from total DNA amplicon libraries of all samples over time faceted by chaparral burned (**E**), woodland burned **(F)**, chaparral post-fire control **(G)** and woodland post-fire control (**H**). Each stacked bar is one sample, and all other parameters are like **C/D**. For **C-H**, sample IDs consist of time point (T1-T5) if applicable, field site (M for McLaughlin or Q for Quail Ridge), habitat (C for chapparal, W for woodland), plot number (1-4), and replicate (A or B). Legend colors are for **C-H**, with the “mixed” category only referring to **C/D**, where predicted hosts came from multiple phyla.

Only 11% of vOTUs were uniquely found in both burned habitats and 1.3% in both control habitats. This signifies that it is likely unique vOTUs that are driving community differences in the burned habitats, rather than shifts in the relative abundances of shared vOTUs (**Supplementary Figure 1A**).

For prokaryotic communities, burn condition explained 12% of the variation, habitat explained 8.2%, and the interaction between burn condition and habitat explained 5.6% (all p < 0.001 by PERMANOVA) (**Figure 2B**). In terms of dispersion, prokaryotic communities in woodland and chaparral were more similar to each other in burned samples than control samples, despite habitat differences (p = 0.002 by PERMDISP). This suggests a homogenizing effect of fire on prokaryotic communities, which was also found to be the case in a previous study of post-wildfire surface soils in a conifer forest [50]. Given the distinct separation between control and burned samples in soil chemical properties, prokaryotic, and viral communities across space, time, and habitat, this wildfire likely had a more homogenous impact on the landscape than did a prescribed burn in our previous study of a forest habitat [39], likely attributable to overall greater fire severity. While heterogenous burn severity over short distances best explained community compositional responses in the previous study, with low-severity burned communities appearing similar to unburned controls and generally only high-severity communities exhibiting a significant response to burn, here the response was more uniform across burned samples.

### Taxonomic shifts in burned versus control samples differed by habitat

To assess whether viral infection dynamics may have changed after wildfire, we next compared viromes according to predicted host taxonomic profiles between burned and unburned samples, leveraging data from T3 (March 2021, approximately five months since the fire), from which we could make these direct comparisons (due to low DNA yields, an insufficient number of viromes was sequenced from other time points). Relative to the woodland habitat, more significant shifts in the relative abundances of viral groups according to predicted host taxonomy were observed for chaparral, which showed a significant increase in the relative abundances of phages predicted to infect Actinobacteria after wildfire and significant decreases in the relative abundances of putative phages of Acidobacteriota and Nitrospirota (Kruskal-Wallis, p < 0.05) (**Figure 2C**). The woodland results were less consistent, with some comparisons showing an increase in putative phages of Actinobacteria, but not uniformly across burned samples. Only the relative abundances of viruses predicted to infect Cyanobacteria had a significant change (Kruskal-Wallis, p < 0.05) between burned and control viromes in the woodland habitat (**Figure 2D**).

We were able to compare the relative abundances of prokaryotic phyla in control and burned samples for all three timepoints from which control samples were collected (T3-T5). In chaparral specifically, multiple prokaryotic phyla remained significantly different in burned samples as compared to control samples until T5 (August 2021, approximately one year since the fire), with some significantly different between burned and control samples across all timepoints (**Figures 2E & G**). The relative abundances of Actinobacteriota and Firmicutes were significantly higher in burned samples at each timepoint, perhaps reflective of the ability of members of these phyla to form spores as survival structures [51,52]. Increased relative abundances of Firmicutes have been previously observed in the first year after a wildfire in chaparral [12] and forest ecosystems [53], as well as in heated soil [9]. Although there was a significant shift in the relative abundances of viruses predicted to infect Nitrospirota, a phylum of nitrite-oxidizing bacteria that typically dominates in arid ecosystems [54], members of this phylum were at very low relative abundance in the prokaryotic data and did not exhibit the same pattern as their viruses. In woodland habitats (**Figures 2F & H**), similar to the patterns in the viruses when grouped by predicted hosts, differences in prokaryotic communities between burned and control samples were less pronounced and were observed at fewer time points.

Firmicutes were significantly more abundant in burned samples (only at T4 and T5), similar to our observations in chaparral (Kruskal-Wallis, p < 0.05). In general, the woodland habitat did not exhibit as pronounced a response to the fire, in terms of phylum-level shifts in the relative abundances of prokaryotes or their predicted phages.

Of the three timepoints that had both burned and unburned samples (T3-T5), only one timepoint had sufficient data to compare viral community compositional shifts between the two sample types. Thus, we were unable to track changes over time in burned and unburned soils. However, we were able to determine whether there was significant compositional changes for viruses (based on host-prediction data) or prokaryotes as the days since fire increased (**Supplementary Figures 1B-E**). In chaparral, there were significant changes in the relative abundance of viruses predicted to infect Actinobacteria and Bacteroidota (decreasing and increasing over time, respectively), while in woodland, putative viruses of Bacteroidota and Proteobacteria significantly decreased over time (Spearman’s, p < 0.05). Viruses predicted to infect all other phyla with significant changes in their relative abundances across timepoints (Kruskal-Wallis, p < 0.05) had trends that were non-monotonic over time. For prokaryotes, most phyla that significantly differed in relative abundance across timepoints (Kruskal-Wallis, p < 0.05) also had non-monotonic relationships with days since fire, potentially reflecting seasonal changes. Only Actinobacteria significantly increased over time in both chaparral and woodland habitats, and Acidobacteriota decreased over time in woodland (Spearman’s, p < 0.05).

#### Biogeographic context suggests that some viral taxa were preferentially detected in post-fire datasets and thus may be fire-adapted

To assess biogeographic patterns of fire-impacted vOTUs, we performed read mapping of this LNU viromic dataset to other vOTU databases to see whether our viromes included vOTUs that had been previously detected elsewhere, including in our other fire-related studies. We detected a total of 1,644 such vOTUs, with 760 from a mixed conifer forest soil as part of our prescribed burn study in Blodgett Forest, CA [9,39] and 884 from the PIGEON v2.0 database [38]. The overall composition of vOTUs in the PIGEON v2.0 database together with the Blodgett Forest vOTUs reflected primarily aqueous (freshwater and marine), forest, and agricultural sources (**Figure 3A**). Other than vOTUs from our November 2019 viromes from these sites, which we had already included in this study (and therefore removed from the database, leaving a total of 787,363 vOTUs), the PIGEON v2.0 database did not contain any vOTUs known to originate from chaparral or woodland habitats, though some vOTUs were from samples labeled as ‘terrestrial (soil)’ in the original publication [55], which could conceivably have included these habitats. When separated by habitat and burn condition, a large percentage of vOTUs detected in burned chaparral (total vOTUs = 527) (**Figure 3B**) and burned woodland (total vOTUs = 912) (**Figure 3C**) viromes were originally found in burned mixed conifer forest (31.3% and 18.9% respectively). Conversely, when considering the composition of vOTUs found in the control viromes (612 total vOTUs for chaparral and 543 total vOTUs for woodland), burned mixed conifer forest was a much smaller fraction (6.3% for chaparral, **Figure 3D**, to almost undetectable at 0.6% for woodland, **Figure 3E**). This suggests that there may be specific groups of viral ‘species’ that commonly persist in post-fire environments, presumably in part due to survival and/or rapid colonization and growth of their hosts [10], but also perhaps due to their own inherent survival traits (e.g., as genomes inside spore-forming hosts and/or as durable virions) [56,57].

**Figure 3.**
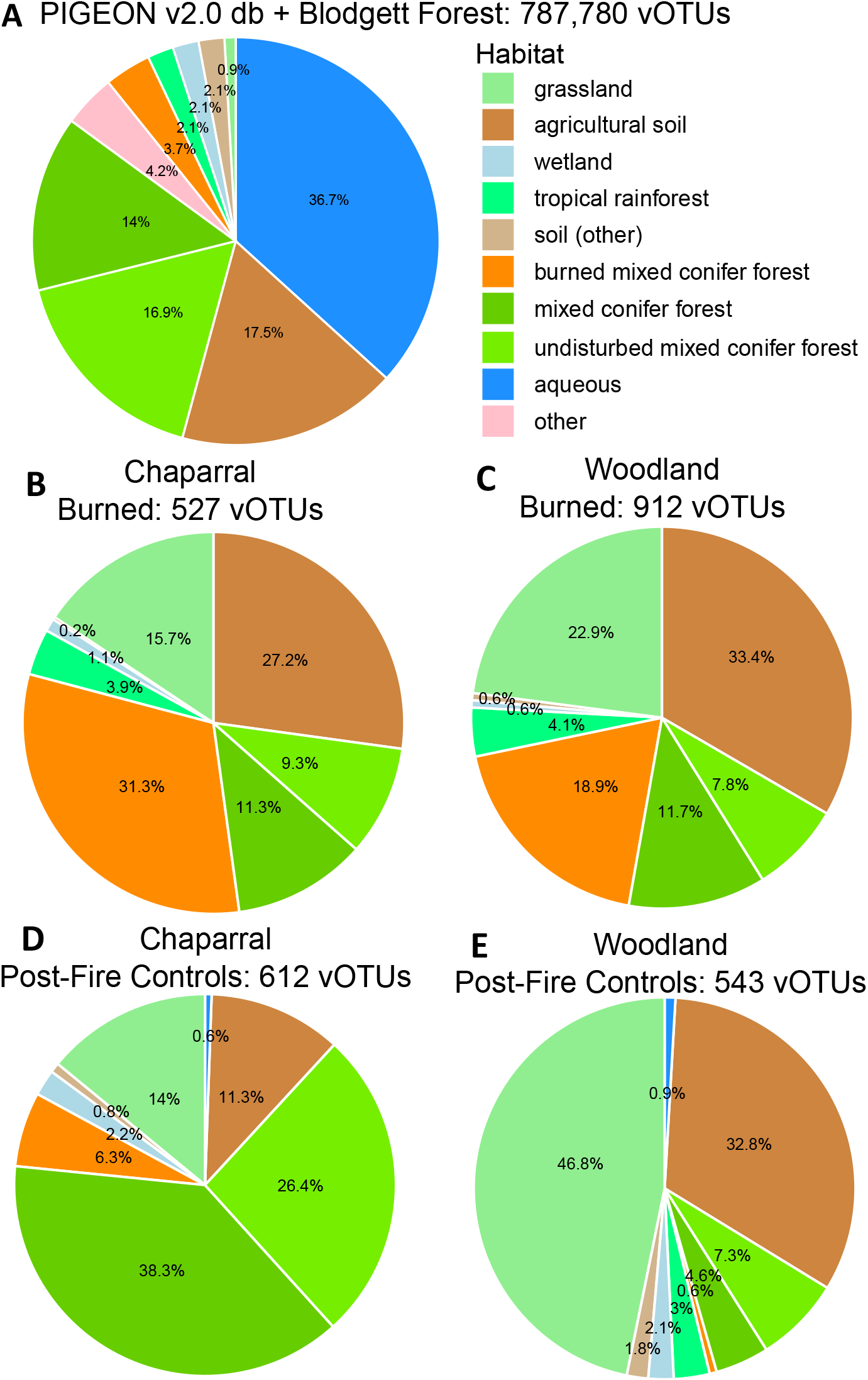
**A**. Habitat composition (percentages) of 787,780 vOTUS from the combined set of vOTUs from the of PIGEON v2.0 database (db) and the Blodgett Forest prescribed burn and laboratory heating studies [9,39], colored by original source environment. **B-E**. Relative proportions of the vOTUs from (**A**) that were detected in our dataset, separated by the habitat (chaparral or woodland) and burn status (burn or control) in which they were detected.

## CONCLUSION

In this study, we compared viral and prokaryotic community responses to wildfire in chaparral and woodland habitats over one year. Many of our viromes had low-to-undetectable viromic DNA yields, suggesting that post-fire conditions in arid climates likely have low extracellular viral biomass. Of the 39 viromes that we were able to sequence, significant differences were observed between burned and control viromes for both habitats, alongside significant changes in soil chemical properties and prokaryotic community composition. Wildfire increased dispersion in viral community beta-diversity for both habitats, while there was an opposite, homogenizing effect on prokaryotic community composition, with prokaryotic communities from different habitats more similar in burned samples than in controls. At T3 (the only timepoint from which control viromes were sequenced), significant increases in the relative abundances of viruses predicted to infect Actinobacteriota and decreases in putative Nitrospirota and Acidobacteriota phages were evident in chaparral, while only viruses predicted to infect Cyanobacteria significantly decreased in woodland. As for prokaryotic shifts, only Actinobacteria and Firmicutes were consistently significantly more abundant in burned chaparral samples at all timepoints than the controls (suggesting increased survival of spore-formers), with no consistent changes in woodland. Additionally, there seemed to be a degree of habitat filtering on viral ‘species’ in burned ecosystems, regardless of habitat, given that a substantial percentage of vOTUs in burned chaparral and woodland were also found previously in burned forest.

In summary, wildfire had a clear impact on viral community composition, including changes in the relative abundances of viruses predicted to infect bacteria from spore-forming phyla, which were at higher abundance in burned samples. Thus, the shifts in viral community composition post-fire were at least partially attributable to shifts their host abundances, but perhaps some of these viruses were also inherently more durable (fire-resistant). Higher temporal resolution in both burned and unburned viromic data will help us to better characterize these compositional shifts over time and compare them with prokaryotic community shifts, enhancing our understanding of virus-host dynamics post-fire.

## DATA AVAILABILITY

All raw sequences have been deposited in the NCBI Sequence Read Archive under the BioProject accession PRJNA1147726. The database of dereplicated vOTUs is available at https://zenodo.org/records/12740558. All scripts are available at https://github.com/seugeo/wildfire.

## ACKNOWLEDGEMENTS

We thank Cathy Koehler at McLaughlin Natural Reserve and Shane Waddell at Quail Ridge Natural Reserve for advising on field sampling logistics and coordinating site visits, and Jane Fudyma, Kristin Kawecki, and David Robles for help with soil collection. Thanks to Valerie Eviner and Jorge Rodrigues for helpful comments on a draft version of the manuscript, Luke Hillary for help with data analysis, and Lucie Jiraska and Matthew Perry for consultation on the abstract. Funding for this work was provided by the U.S. Department of Energy (DOE), Office of Science, Office of Biological and Environmental Research (BER), Genomic Science Program, award number DE-SC0021198 (grant to JBE). SEG was also supported by the National Science Foundation Graduate Research Fellowship, the University of California Eugene Cota-Robles Fellowship, and the College of Agriculture and Environmental Sciences (UC Davis) Dean’s Circle Award.

---

**Supplementary Figure 1.**
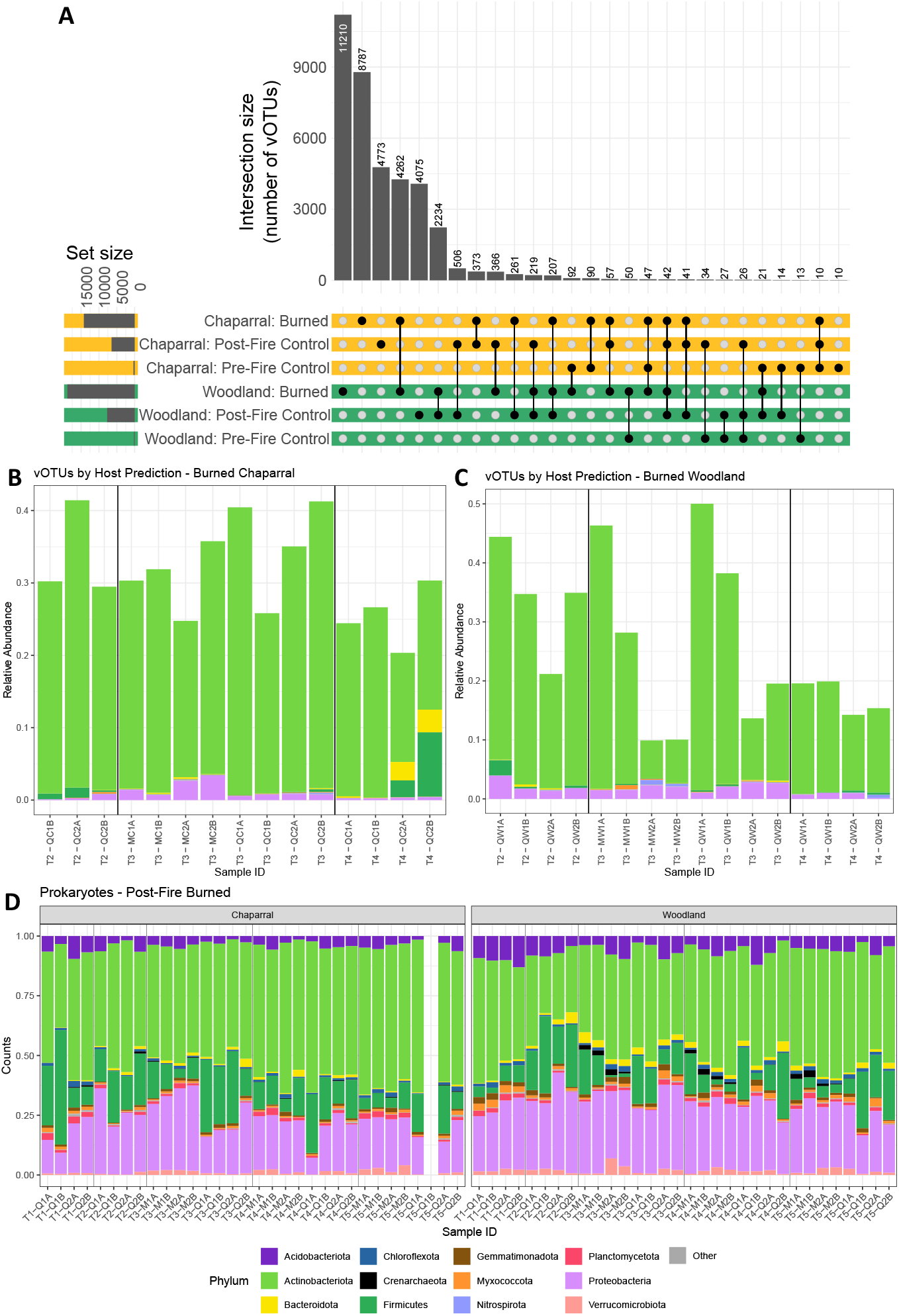
**A**. UpSet plot of vOTU detection patterns across plots with blackened circles indicating detection in a particular habitat with a particular condition (burned, pre-fire control, and post-fire control) and intersection size indicating the number of vOTUs with a given detection pattern (detected in one or more habitat/burn condition combinations). **B/C**. Time series showing the relative abundances of vOTUs grouped by their phylum-level host predictions for burned chaparral (**B**) and burned woodland (**C**) time points. Each bar is a virome, and vertical black lines separate time points. All lower abundance vOTU groups (according to host phyla) are collapsed into an “other” group. **D**. Time series showing relative abundances of prokaryotic phyla, faceted by habitat (chaparral – left, woodland – right) from burned samples from all timepoints. Each bar is a sample, with the missing bar representing a sample that failed amplification and was not sequenced. Vertical black lines separate time points. All lower abundance phyla are collapsed into an “other” group. For **B-D**, sample IDs consist of time point (T1-T5), field site (M for McLaughlin or Q for Quail Ridge), habitat (C for chapparal, W for woodland), plot number (1-4), and replicate (A or B).

